# Torpor duration, more than temperature, is key to hummingbird nighttime energy savings

**DOI:** 10.1101/383679

**Authors:** A Shankar, RJ Schroeder, SM Wethington, CH Graham, DR Powers

**Author notes:** Equal and corresponding authors.

## Abstract

1. Torpor is an important energy saving strategy in small endotherms, but it has been insufficiently studied in natural field conditions. Building on what we know from laboratory studies, we compared torpor use across hummingbird species and different natural temperature regimes to explore predominant hypotheses about torpor use and evaluate the possible effects of environmental variation on energy management.
2. We found that the probability of an individual entering torpor was correlated with mass and unrelated to nighttime temperature, and that hummingbirds at both warm tropical and cooler temperate sites used torpor.
3. Energy savings in torpor were maximized as ambient temperatures approached a species’ minimum body temperature consistent with laboratory studies; energy savings ranged between 65-92% of energy per hour in torpor compared to normothermy.
4. However, variation in total nighttime energy expenditure was more significantly influenced by torpor bout duration than by the variation in energy savings in torpor.
5. Our results show that a small endotherm’s nighttime energy management in its natural habitat is more affected by torpor bout duration, which is linked to photoperiod, than by temperature. This result suggests that in their natural environments, hummingbirds are able to save energy in torpor across a range nighttime temperature, indicating that they may have sufficient physiological flexibility to tolerate climatic variation.

## INTRODUCTION

Torpor is an important strategy for energy management in over 200 endotherm species (Ruf & Geiser, 2014). In torpor, an animal lowers its body temperature (T_b_) down to a minimum T_b_ (sub-zero to 30°C) as ambient temperature (T_a_) decreases (Geiser, Currie, Shea, & Hiebert, 2014; Kruger, Prinzinger, & Schuchmann, 1982; Reinertsen, 1996; Richter et al., 2015a). Torpor allows an animal to save energy by eliminating the costs of regulated heat production (Hainsworth & Wolf, 1970; Ruf & Geiser, 2014; Fig. 1). Controlled laboratory studies indicate that daily torpor use in small endotherms is influenced by environmental factors, such as T_a_. These studies provide significant insight into the energy savings gained from torpor, but they generally focus on a single environmental factor; in the wild multiple environmental factors can vary simultaneously (Boyles, Seebacher, Smit, & McKechnie, 2011). Further, the influence of natural energy use and storage throughout the day is not considered, though it can greatly impact torpor use (Powers, Brown, & Van Hook, 2003). As a result, laboratory studies may not always provide a comprehensive picture of torpor under natural conditions, highlighting the need for field physiology studies. Species often experience a range of environmental conditions, and therefore, it is important to evaluate their ability to adjust to a variety of conditions, such as variable temperature, photoperiod, and resource availability. Here, we evaluate torpor use in eight species of small endotherms, hummingbirds, and assess the factors influencing nighttime energy savings under natural temperatures and photoperiods at both temperate and tropical sites.

**Figure 1:**
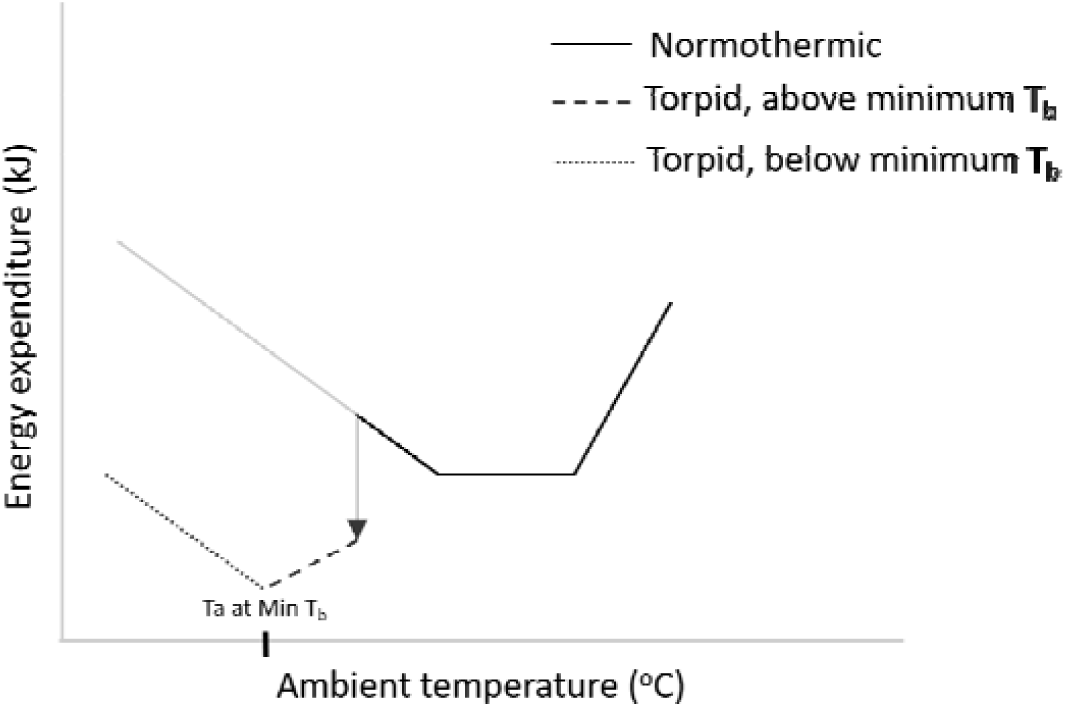
Theoretical relationship between energy expenditure and ambient temperature. The solid lines depict normothermic parts of the Scholander-Irving curve (Scholander et al. 1950), and dashed lines represent torpid regions (adapted from data in (Hainsworth & Wolf, 1970) and this study). Energy expenditure in torpor can drop down until minimum body temperature (min T_b_) is reached, after which energy must be expended (dotted line) to maintain min T_b_.

Hummingbirds, which are among the smallest vertebrate endotherms, are recognized for their ability to use daily torpor to reduce their nighttime energy expenditure (Kruger et al., 1982; NEE; Pearson, 1950; Schuchmann, Kruger, & Prinzinger, 1983). Laboratory studies have provided a clear understanding of the individual relationships between torpor use and the depression of T_b_, the metabolic defence of minimum T_b_ (Carpenter, 1974; Hainsworth & Wolf, 1970; Wolf & Hainsworth, 1972), and endogenous energy stores (Hiebert, 1992; Powers et al., 2003). Laboratory measurements of metabolic rates in torpor indicate that torpid hummingbirds maximize energy savings when T_a_ – T_b_ is minimized and they are at their minimum T_b_ (Hainsworth & Wolf, 1970; Jastroch et al., 2016). Minimum T_b_ varies by species (4-18°C, Hainsworth and Wolf 1970, Carpenter 1974), suggesting that it has an adaptive role; however the importance of minimum T_b_ for hummingbird energy management under natural conditions has not been assessed. Because laboratory studies use controlled temperature steps to evaluate the impact of temperature on torpor metabolism (Kruger et al., 1982; Lasiewski, 1963; Powers et al., 2003), patterns of T_b_ management might not reflect torpor use under natural T_a_ cycles, which are generally more predictable because T_a_ tends to decline gradually through the night. In addition, laboratory measurements do not always incorporate the natural patterns of energy storage in free-living birds (Auer, Bassar, Salin, & Metcalfe, 2016), which can affect whether or not torpor is used (Hainsworth, Collins, & Wolf, 1977; Powers et al., 2003). Laboratory measurements of torpor can also underestimate torpor bout duration and reduction of T_b_ (Geiser et al., 2000). A few studies have measured torpor while incorporating some of the natural conditions hummingbirds experience. For instance, body temperatures of hummingbirds in torpor under natural temperature cycles have been measured, but the influence of natural energy stores and photoperiod on torpor were not considered (Bech, Abe, Steffensen, Berger, & Bicudo, 1997). Torpor use has also been measured on hummingbirds that naturally acquired energy stores, but the influence of photoperiod or temperature cycles were not simultaneously evaluated (Powers et al., 2003). Thus, it is unknown whether hummingbirds maximize energy savings by maximizing torpor bout duration or if maximizing energy savings occurs when T_a_ approaches minimum T_b_, or both.

Three major hypotheses have thus far been used to explain patterns of hummingbird torpor use. Under the emergency-only hypothesis (Hainsworth et al., 1977), hummingbirds use torpor only when energy stores are below some critical threshold. This threshold is usually low, though it may be higher during migration when birds maximize fat gain (Carpenter & Hixon, 1988). Under the routine hypothesis (Kruger et al., 1982), hummingbirds use torpor every night, while under the circadian rhythm hypothesis (Carpenter, 1974), torpor is an innate circadian response to day length and season. The relative importance of these three hypotheses might be influenced by food availability, mass, degree of territoriality and T_a_. First, sufficient food is required to maintain an energy buffer to survive the night and support morning foraging, as hummingbirds appear to have an endogenous energy storage threshold that triggers torpor when crop storage and fat are depleted (Bicudo, 1996; Calder, 1994; Hiebert, 1992; Powers et al., 2003). Second, a species’ degree of territoriality and size can be linked to its fat storage patterns: the territorial blue-throated hummingbird (*Lampornis clemenciae*, ca. 8g), appeared to only use torpor in emergencies, because it actively defends resources and can access and store energy for nighttime use (Powers et al., 2003). In contrast, non-territorial species in similar environments do not have easy access to resources and might use torpor more routinely (Hainsworth, 1978; Hainsworth et al., 1977; King, 1970; Powers & Wethington, 1999). For example, non-territorial Rivoli’s hummingbirds (*Eugenes fulgens*, ca. 7.5g) and black-chinned hummingbirds (*Archilocus alexandri*, ca. 3g) enter torpor regularly (Calder, 1994; Powers et al., 2003).

We evaluated patterns of torpor use in free-living hummingbirds that naturally acquired energy through the day at several sites under natural nighttime temperatures and photoperiods. We also evaluated the role of minimum T_b_ and torpor duration in total nighttime energy savings, asking whether nighttime energy savings (by lowering T_b_) and duration differed between sites with shorter, cooler nights vs. those with longer, warmer nights. We predicted that as nighttime temperatures got colder or hotter than minimum T_b_ and hourly energy savings in torpor would decrease and total nighttime energy expenditure would be higher. Thus, we predicted that temperature would have the largest effect on nighttime energy expenditure. We tested these predictions in eight hummingbird species across five sites in North and South America.

## METHODS

### Study sites and species

We collected data at five sites: three in Arizona, USA and two on the eastern slope of the western Ecuadorian Andes. We studied eight species, and used only males to avoid potential disturbances to nesting (summarized in Table 1).

**Table 1:** Characteristics of the study sites - three temperate (in Arizona, AZ) and two tropical sites (in Ecuador, EC).

| Region | Site | Species | Territorial (T)/<br>Non-territorial<br>(NT) | Latitude,<br>Longitude | Altitude<br>(m) | Dates<br>sampled | Hummingbird-<br>visited plant<br>species | Night<br>length<br>(hours) |
| --- | --- | --- | --- | --- | --- | --- | --- | --- |
| AZ | Harshaw<br>Creek | *Cynanthus latirostris* (CYLA) | NT | 31.50,<br>-110.68 | 1370 –<br>1635 | Jun – Jul<br>2013 | 11 | 9 |
| AZ | Sonoita<br>Creek | *C. latirostris* (CYLA) | NT | 31.50,<br>-110.86 | 1100 –<br>1180 | Jun – Jul<br>2013 | 7 |  |
| AZ | Southwestern<br>Research<br>Station<br>(SWRS) | *Eugenes fulgens* (EUFU)<br>*Lampornis clemenciae* (LACL) | NT<br>T | 31.88,<br>-109.21 | 1650 –<br>1720 | May – Jul<br>2014 | 29 | 11 |
| EC | Maquipucuna | *Phaethornis syrmatophorus* (PHSY)<br>*Heliodoxa jacula* (HEJA)<br>*Florisuga mellivora* (FLME) | NT<br>T<br>T | 0.12,<br>-78.64 | 1275 –<br>1370 | Jul – Aug<br>2014 | 30 |  |
| EC | Santa Lucia | HEJA<br>FLME<br>*H. imperatrix* (HEIM)<br>*H. rubinoides* (HERU) | T<br>NT<br>T<br>T | 0.12,<br>-78.61 | 1800 –<br>2100 | Jul – Aug<br>2014 | 45 |  |

### Temperature

We measured site-level ambient temperature (T_a_) using temperature loggers (iButtons and Hobo H8 temperature loggers) placed in Styrofoam or plastic containers 1 meter from the ground to insulate them from solar radiation and wind. We measured operative temperature (T_e_), the temperature as experienced by a hummingbird, using copper sphere placed in the environment where a hummingbird would perch (Walsberg & Weathers, 1986). We also measured internal temperature of the metabolism chambers (T_c_) to the nearest 0.1°C continuously during torpor measurements. using either thermocouples or thermistor probes We used T_c_ to check how closely temperatures experienced by the birds in the chamber approximated T_a_ and T_e_ (temperature measurements detailed in Supplement S1).

### Torpor measurement

We used modified Russell traps and mist nets to capture birds about a half hour before dark to allow for near-natural daytime activities and feeding regimens for storage of endogenous energy. Because hummingbirds likely depend, at least in part, on crop sugar for a portion of their nighttime energy expenditure, we allowed birds to feed in cages *ad libitum* for 30 minutes to match excessive evening foraging or hyperphagia (Calder et al. 1990; Powers et al. 2003), prior to being placed into metabolism chambers (AZ 10.0 litre, 23.5 × 19.5 × 22.0 cm; Ecuador 6.0 litre, 26 × 18 × 15 cm). We used negative-pressure open-flow respirometry to measure nighttime metabolism and track the use of nocturnal torpor (see Supplement S1: Figs. S1, S2 and associated text for respirometry methods; Powers et al. 2003, Welch 2011). Even though we used natural temperature patterns and photoperiods, we cannot rule out the possible effects of capture, handling and behavioural differences between species, which could affect patterns of torpor use in these birds. However, we believe that our methods better approximate what hummingbirds experience in the wild than other current protocols. We assumed birds entered into torpor when metabolic rate fell below resting normothermic values (a minimum of 0.4 O2 mL/min change, and an average of 1.1 mL O2/min in 30-90 minutes; Hiebert, 1990; Powers et al., 2003). We terminated measurements about 30 minutes after rewarming ended but no earlier than 30 minutes prior to sunrise (Calder, 1975) and allowed birds to feed in holding cages for 30 minutes prior to release. We calculated rewarming costs by integrating the area under the curve starting when VO_2_ began to steadily increase and ending when the birds stopped actively increasing their metabolism (Bartholomew & Lighton, 1986). For broad-billed hummingbirds (n=15), we additionally measured energy expenditure under controlled laboratory conditions from 5-30°C in 5°C increments to compare to field data and estimate minimum T_b_ (Supplement S1).

### Torpor analyses

Torpor data were processed using Warthog Lab Analyst software (warthog.ucr.edu). We obtained VO_2_ values for analysis during times when the bird was in equilibrium. Our VO_2_ measurements were converted to energy expended in kJ/hour and summed over the night to find NEE (kJ). To convert VO_2_ to energy units, we assumed 20.1 J/ml O_2_ (Kleiber, 1975).

A standardized set of measures to compare torpor use between species are lacking in the literature, though these measures have previously been used independently. We calculated the following four measures to evaluate torpor use.

1. *Frequency of torpor* is the percentage of individuals that entered torpor per species (Powers et al., 2003).
2. *Duration* is the average number of hours (to the nearest half-hour) a species used torpor per night, including torpor entry and excluding rewarming. To accommodate variation in available night length, we also report the percentage of nighttime hours spent in torpor.
3. *Hourly energy savings* is the percentage of energy saved in torpor relative to normothermy for an individual, and is useful in comparing across sites with variable night lengths. This measure has also been referred to as “relative torpor metabolic rate” (Ruf & Geiser, 2014) and percentage energy expenditures (Bucher & Chappell, 1997; Schuchmann et al., 1983). Torpor and normothermy values were obtained by averaging hourly torpid and normothermic energy expenditure over the entire night for each individual. Note that all individuals spent at least part of the night normothermic (i.e., before torpor entry and after exiting) allowing us to obtain normothermic values for torpid individuals (Supplement S1 for calculation details).

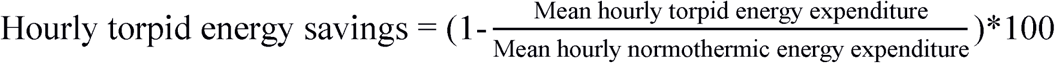
4. *Total nighttime energy expenditure* is per individual mass-corrected nighttime energy expenditure (NEE/mass; Lighton, 2008).

### Bird mass

We weighed each individual three times to obtain a capture mass, a fed mass before placing the bird in the chamber, and an end mass when the bird was removed the next morning. We used the capture mass in subsequent analyses (see Supplement S1 for details of mass measurements, Table S1 for fractional mass change).

### Statistical analyses

We used the phylogeny generated by McGuire et al. (2014), pruned to match our dataset, to perform phylogenetically corrected analyses that accommodated the lack of independence caused by the species’ relatedness. We used a hierarchical Bayesian approach to accommodate the hierarchical nature of individual observations clustered within species. We ran phylogenetic generalized linear mixed models using a Markov chain Monte Carlo (MCMC) sampler in the R package MCMCglmm v. 2.24 (Hadfield, 2010) which models the phylogenetic relatedness between species as a random variable. We ran the MCMC chain for 5 million iterations, sampling every 1,000 generations. We used uninformative priors, fixing the variance for the residual to 1 (see Supplement S1 for modelling details).

Using this Bayesian hierarchical MCMCglmm framework we first assessed whether the probability of entry into torpor (modelled as an individual-level binary variable) was affected by individual capture mass. We then evaluated which factors (mass, torpor bout duration, minimum T_c_, rewarming costs, and hourly energy savings) best predicted total mass-corrected nighttime energy expenditure considering both the individual variables and a forward stepwise model. We compared these models using their DIC scores. Finally, we tested whether energy expenditure during rewarming was correlated with mass and T_c_ during rewarming by comparing DIC outputs from mass-only and mass + T_c_ models. All the MCMCglmm results are reported as post-mean with credible intervals (CI) and pMCMC; if the reported CI’s do not overlap zero, we infer that variable does influence the structure of the data (e.g., if the post-mean and credible intervals are negative, that variable has a negative effect on the dependent variable and vice versa; Hadfield, 2010). We also used t-tests to analyse differences in torpor bout duration, hourly energy savings, and mass-corrected nighttime energy expenditure between individuals at temperate and tropical sites, as well as between sites for which we had sufficient sample sizes. All t-tests were unpaired two-tailed t-tests with equal variances, since all groups tested had equal variances.

## RESULTS

### Temperature

Metabolic chamber temperature (T_c_; Fig. 2) generally tracked ambient temperature (T_a_; Supplement S1: Fig. S3), indicating that our chambers reflected natural temperature patterns. The only exception was Harshaw, where T_a_ had greater variability through the night than T_c_, because the habitat and topographic complexity at this site was not reflected in the chambers. Nighttime T_c_ were generally warmer at tropical sites, more variable at temperate sites and declined through the night at all sites.

**Figure 2:**
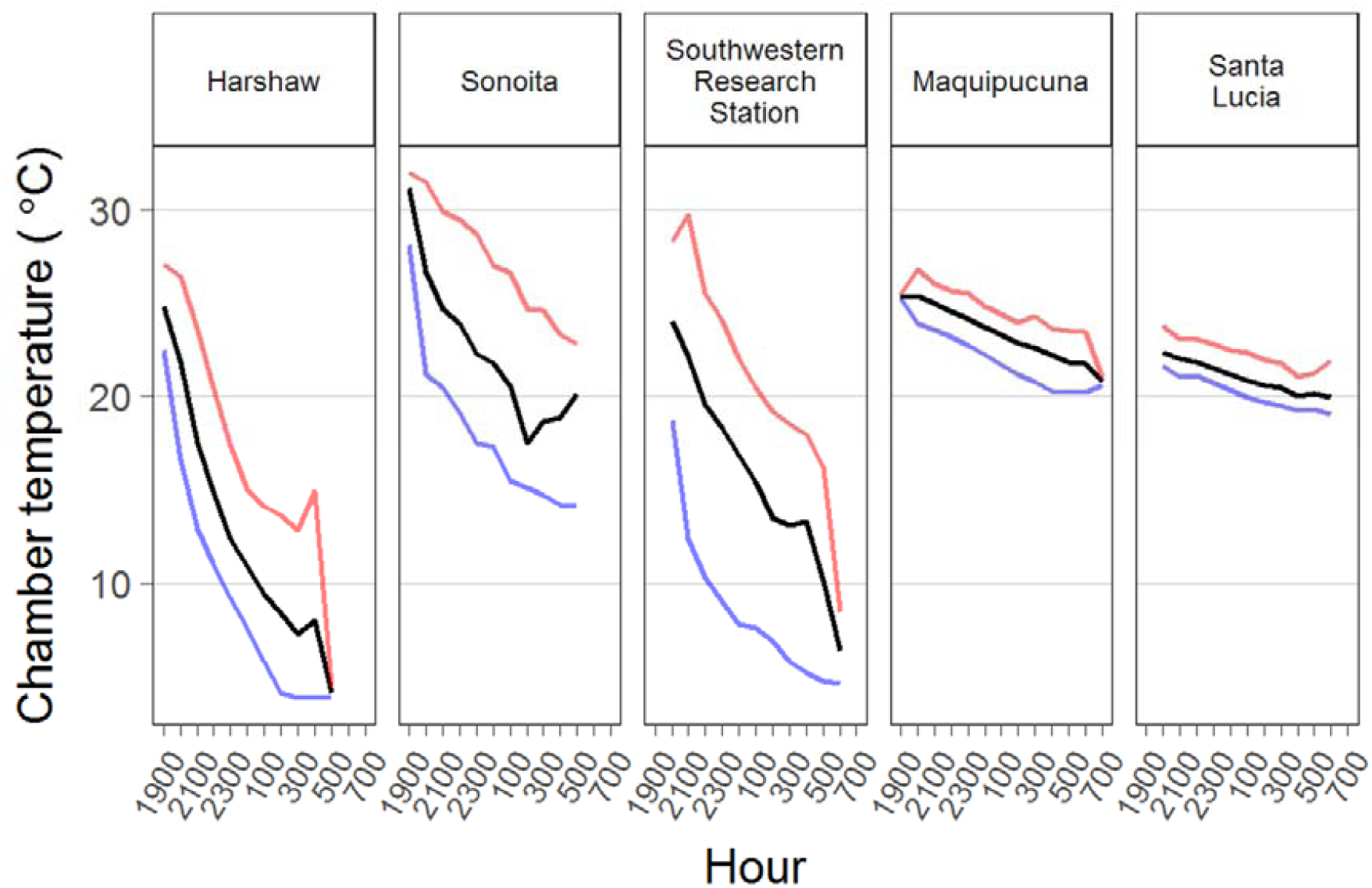
Chamber temperatures averaged per hour from all experiment nights, across sites. The dark lines denote the average across days at that hour of night; lighter lines denote maximum and minimum chamber temperatures at that hour. The first three facets are temperate sites, the last two facets are tropical sites.

### Frequency and probability of entry into torpor

The overall frequency of torpor for all individuals studied was 54.8% (23 torpid/42 birds; see Table 2 for species-specific details). Two of the territorialist species—the blue-throated hummingbird and the empress brilliant—never entered torpor, while all other species entered torpor with at least 25% frequency. Individuals exposed to the same environmental conditions varied in whether they used torpor. For example, at the Southwestern Research Station, we twice measured two Rivoli’s hummingbirds on the same night and both times one bird used torpor while the other did not. The MCMCglmm model for the probability of an individual entering torpor as a function of capture mass indicated that heavier birds had a lower chance of entering torpor (Table 3). This model estimated a positive mean intercept and negative mean slope (Table 3; Supplement S1: Fig. S4).

**Table 2:** Measures of nighttime energy expenditure and torpor use per species at a site, presented as normothermic vs. torpid birds. *SWRS = Southwestern Research Station

| Site | Species | N vs. T | Mean capture mass (g) | Mean (range) mass-corrected NEE (kJ/g) | Probability of entering torpor | Mean (range) torpor duration (hours) | Mean hourly torpid energy savings |
| --- | --- | --- | --- | --- | --- | --- | --- |
| Harshaw | *CYLA* | 0 vs. 7 | 3.20 | NA vs. 1.05 (0.73 – 1.52) | 1 | 4.7 (2.5 – 6) | 82 % |
| Sonoita | *CYLA* | 3 vs. 5 | 3.34 vs. 3.23 | 1.73 (1.35 – 2.02) vs. 0.96 (0.65 – 1.24) | 0.625 | 4.1 (2.5 – 7) | 90 % |
| SWRS* | *LACL* | 4 vs. 0 | 8.28 | 2.58 (2.13 – 3.26) vs. NA | 0 | NA | NA |
| SWRS* | *EUFU* | 2 vs. 2 | 7.93, vs. 7.75 | 2.71 (2.68 – 2.75) vs. 2.43 (2.31 – 2.55) | 0.5 | 3.25 (2 – 4.5) | 85% |
| Maquipucuna | *HEJA* | 1 vs. 6 | 8.74 vs. 8.64 | 2.00 vs. 1.01 (0.66 – 1.49) | 0.83 | 6.8 (3 – 8) | 75% |
| Maquipucuna | *PHSY* | 0 vs. 1 | 5.24 | NA vs. 2.06 | 1 | 5 | 78% |
| Maquipucuna | *FLME* | 0 vs. 1 | 6.93 | NA vs. 1.78 | 1 | 3 | 65% |
| Santa Lucia | *HEIM* | 5 vs. 0 | 9.60 | 1.92 (1.23 – 2.19) vs. NA | 0 | NA | NA |
| Santa Lucia | *HERU* | 3 vs. 1 | 8.27 vs. 7.89 | 2.21 (2.08 – 2.29) vs. 1.31 | 0.25 | 4 | 81% |
| Santa Lucia | *HEJA* | 0 vs. 1 | 8.74 | NA vs. 0.86 | 1 | 7 | 89% |
| Santa Lucia | *FLME* | 1 vs. 0 | 6.90 | 1.94 vs. NA | 0 | NA | NA |

**Table 3:**
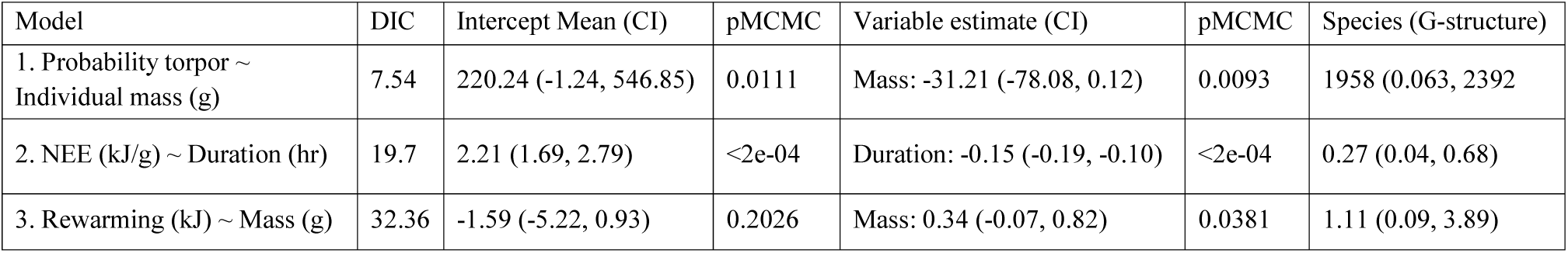
The results of hierarchical Bayesian MCMCglmm models, where species relatedness was accounted for: 1. tests the influence of mass on the probability of entry into torpor, 2. tests the factors influencing mass-corrected nighttime energy expenditure (NEE), 3. tests the extent to which mass influenced rewarming energy expenditure.

| Model | DIC | Intercept Mean (CI) | pMCMC | Variable estimate (CI) | pMCMC | Species (G-structure) |
| --- | --- | --- | --- | --- | --- | --- |
| 1. Probability torpor ~ Individual mass (g) | 7.54 | 220.24 (-1.24, 546.85) | 0.0111 | Mass: -31.21 (-78.08, 0.12) | 0.0093 | 1958 (0.063, 2392) |
| 2. NEE (kJ/g) ~ Duration (hr) | 19.7 | 2.21 (1.69, 2.79) | <2e-04 | Duration: -0.15 (-0.19, -0.10) | <2e-04 | 0.27 (0.04, 0.68) |
| 3. Rewarming (kJ) ~ Mass (g) | 32.36 | -1.59 (-5.22, 0.93) | 0.2026 | Mass: 0.34 (-0.07, 0.82) | 0.0381 | 1.11 (0.09, 3.89) |

### Duration

Torpor bout duration for all individuals that entered torpor was on average 4.9 (range 2.5 – 8) hours (Table 2). Individuals at the tropical sites had marginally higher torpor bout duration (5.9, range 3 – 8 hours) than species at the temperate (4.3, range 2.5 – 7 hours) sites (*t*(14) = −2.09, p = 0.06). There was no difference between temperate and tropical site individuals in the proportion of the night they spent torpid (*t*(17) = −0.70, p = 0.49). Torpor bout duration was unrelated to hourly energy savings and to minimum nighttime T_c_ (Supplement S1: Fig. S9). Individuals that entered torpor earlier in the night had longer torpor bout durations, and all individuals that used torpor had a single, uninterrupted torpor bout (Supplement S1: Figs. S5, S6).

### Torpor energy savings

Average hourly energy savings in torpor across all species was 82% of normothermic costs (Table 2 and Supplement S1: Fig. S7). Hourly torpid energy savings were significantly higher at temperate compared to tropical sites (85% average vs. 76% average; *t*(13) = 2.94, p = 0.01). Average hourly torpid energy savings at Harshaw (Arizona) for broad-billed hummingbirds were lower than hourly torpid savings at Sonoita (Arizona; *t*(10) = 3.07, p = 0.01). Torpor measurements under both semi-natural (natural nighttime temperature gradients) and controlled laboratory conditions (5°C temperature steps) with broad-billed hummingbirds showed similar trends, though birds under natural temperature conditions showed higher variability in energy expenditure, especially at lower temperatures (Fig. 3). Nighttime metabolic rate decreased with decreasing T_c_ as broad-billed hummingbirds reached their minimum tolerable T_b_ (∼15 °C by extrapolating the lower slope at the T_a_ at which the birds started thermoregulating, Fig. 3a, Supplement S1: Fig. S8a); metabolic rate then increased below minimum T_b_.

**Figure 3:**
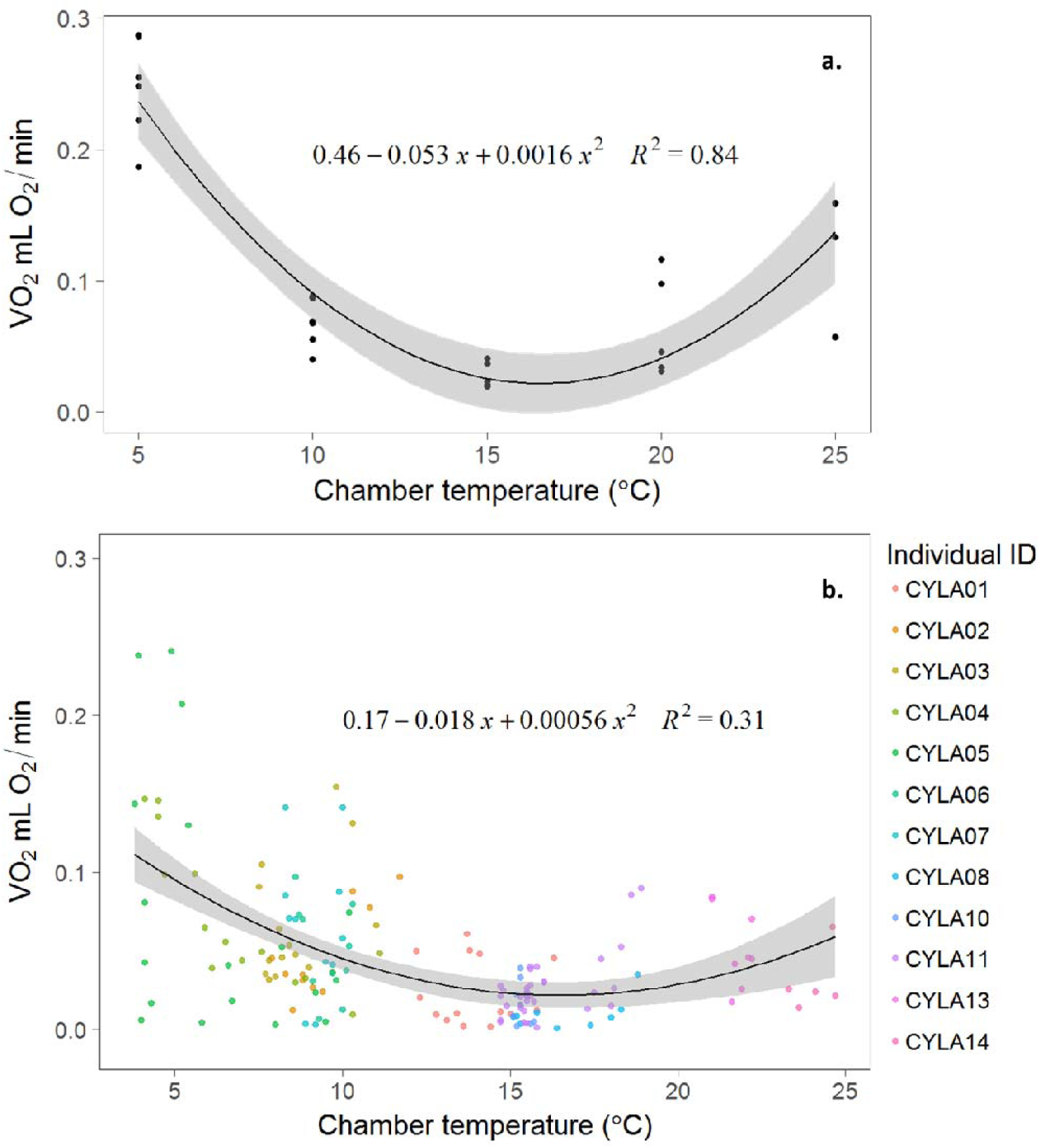
Lab vs. field data on broad-billed hummingbird torpor and oxygen consumption in torpor for broad-billed hummingbirds. a. Oxygen consumption was measured in controlled 5°C temperature steps in a chamber in the lab (5.7 × 15.6 × 5.7 cm) using a Peltier device (Pelt-3; Sable Systems). By extrapolating the lower slope to the x-axis, minimum body temperature for this species seems to be around 15°C. b. Data collected on broad-billed hummingbirds under natural temperature and light cycles in the field. Colours represent individuals. Birds CYLA01 – CYLA07 were measured at the colder site, Harshaw. CYLA08 – CYLA15 were measured at the warmer site, Sonoita. Some of the birds at Sonoita (CYLA09, 12, 15) did not enter torpor and so are not on this graph.

### Nighttime energy expenditure

Mass-corrected total nighttime energy expenditure did not significantly differ between individuals at temperate vs. tropical sites (*t*(39) = 0.04, p = 0.97). Mean, mass-corrected nighttime energy expenditure for torpid broad-billed hummingbirds was not significantly different between Harshaw and Sonoita (*t*(12) = −0.96, p = 0.36). At Sonoita, the total nighttime energy expenditure of normothermic birds was almost double the total nighttime energy of torpid Sonoita birds. Rewarming was more energetically expensive at the colder site, Harshaw, than at Sonoita (*t*(7) = 3.49, p = 0.009), confirming that T_c_ affects rewarming costs within the broad-billed hummingbirds. In Ecuador, mean mass-corrected nighttime energy expenditure overall was marginally lower at Maquipucuna compared to Santa Lucia (*t*(14) = −2.01, p = 0.06). Torpid birds at Maquipucuna had similar total mass-corrected nighttime energy expenditure to torpid birds at Santa Lucia (*t*(3) = 0.61, p = 0.58, Table 2).

The best MCMCglmm models explaining nighttime energy expenditure all included torpor bout duration. Energy savings was significantly negatively related to nighttime energy expenditure in a model including just energy savings; however in both the full and best models energy savings was not significant—only duration was significant. We present the results of the best and most parsimonious model—the duration model—here, and the remaining models in the Supplement S1: Table S2. The duration model showed that mass-corrected nighttime energy expenditure in torpor was negatively related to torpor bout duration, and estimated a positive mean intercept and negative mean slope for duration (Table 3, Supplement S1: Fig. S10). The best and most parsimonious rewarming model included only capture mass, and showed that rewarming costs were positively correlated to individual mass, with a negative mean intercept, and positive mean slope of 0.34 (Table 3, Supplement S1: Figs. S11 and S12). Rewarming duration was an average of 38 minutes (range 7-92 min). Rewarming duration was higher than has been reported for hummingbirds in the past (Hiebert, 1990).

## DISCUSSION

To evaluate existing hypotheses for hummingbird torpor and predict how free-living hummingbirds might adjust their torpor use in response to variable environmental conditions, we assessed the probability of an individual entering torpor and evaluated the effect of torpor bout duration, hourly energy savings, and temperature, on nighttime energy expenditure for hummingbirds across multiple tropical and temperate sites. Hourly energy savings were maximized when the difference between T_a_ and a bird’s minimum T_b_ were close to zero (see Figs. 1 and 4), reflecting what laboratory studies have found in hummingbirds (Hainsworth & Wolf, 1970) as well as in torpid and hibernating mammals (Geiser & Kenagy, 1988; Richter et al., 2015b). Total nighttime energy expenditure was negatively related to hourly energy savings, but our best model for nighttime energy expenditure showed that torpor bout duration was the only factor that caused significant differences in total nighttime energy expenditure. The longer a hummingbird is torpid, the lower its total nighttime energy expenditure, regardless of hourly torpid energy savings. Across all our study sites, including those in the tropics, temperatures were cold enough to allow individuals to enter and use torpor, indicating that temperature is likely not the primary factor determining torpor use. We found that nighttime energy management for hummingbirds likely depends on whether their nights are long enough to use torpor effectively.

**Figure 4:**
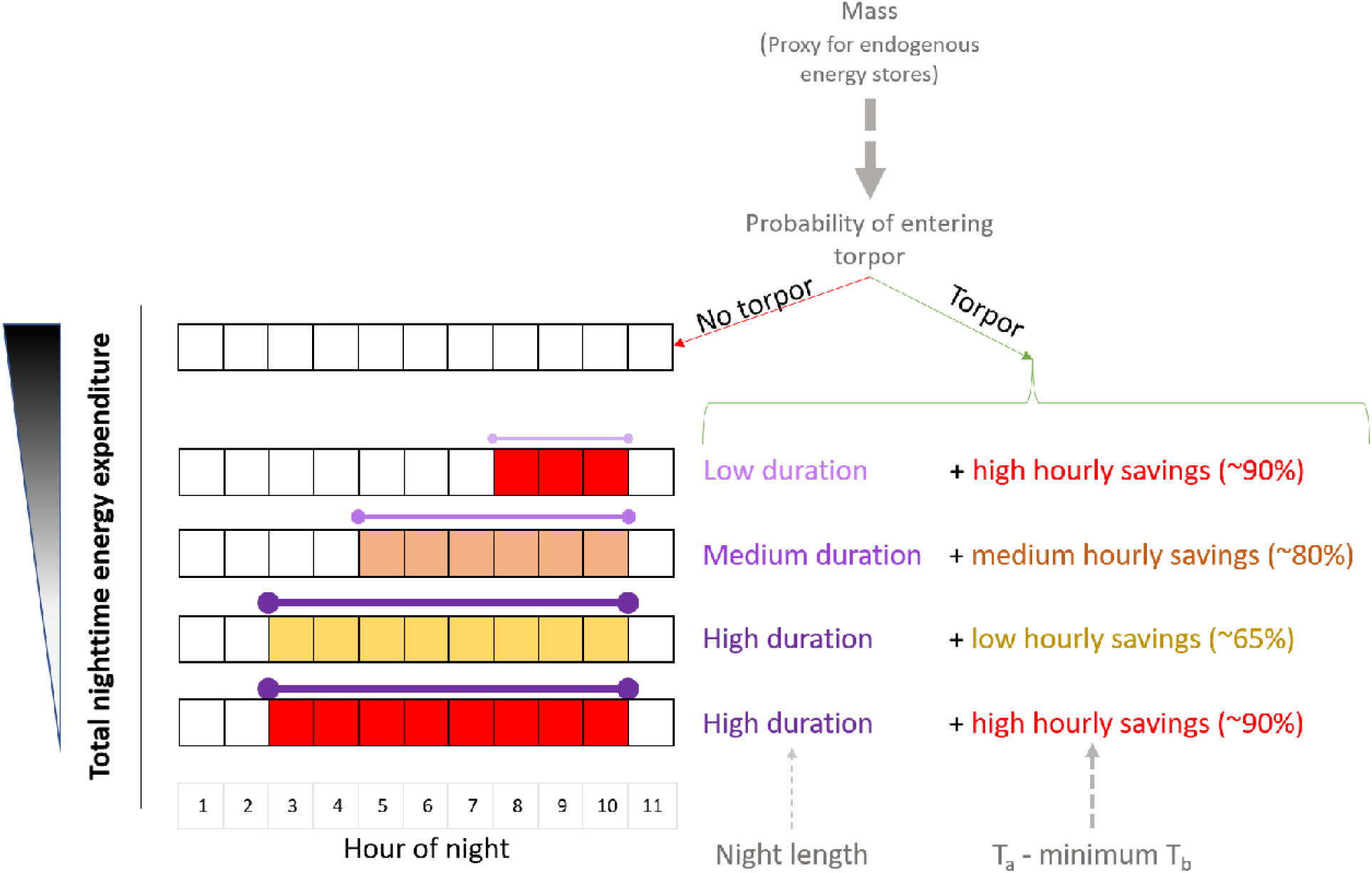
A schematic depiction of aspects of nighttime energy expenditure within a hypothetical hummingbird species. The most important aspects of nighttime energy expenditure (black triangle on left) seem to be the probability of entering torpor (influenced by mass), and th duration of torpor (influenced perhaps by night length), rather than energy savings (which increase as ambient temperature approaches minimum body temperature). A situation with high torpor duration, low hourly energy savings results in lower total nighttime energy expenditure than low torpor duration with high hourly energy savings. Arrows represent the strength of th effect of a variable on a component of nighttime energy expenditure. Dashed arrows represent negative correlations; width of the arrows depict the strength of the correlation.

We predicted that temperature would be the main factor determining the probability of an individual entering torpor; instead, mass was the main factor – heavier species had a lower probability of entering torpor. Heavier, larger hummingbird species have two energetic advantages. First, hummingbirds have limited energy stores for nighttime use because they tend to maximize food intake and energy storage only in the late evening before they roost rather than throughout the day (Calder, Calder, & Frazier, 1990; Powers et al., 2003). Heavier birds can store more energy than smaller birds, either in their crop or as body fat, allowing them to avoid torpor and its associated risks. Second, since metabolic rates have a scaling exponent with body size that is <1 (Glazier, 2015), larger bird species should be able to use normothermy for longer from proportionally similar initial energy stores. In addition to body size, a bird’s capacity for energy storage may be influenced by resource availability and degree of territoriality. The only species to always avoid torpor in our study were two large territorialist species (the empress brilliant and the blue-throated hummingbird). In contrast, species that used torpor consisted of a mix of roles, including smaller territorialists and all non-territorialists. These results suggest that perhaps a combination of the emergency only (Hainsworth et al., 1977; Powers et al., 2003) and routine hypothesis (Kruger et al., 1982) are at play. Large, territorialist hummingbirds might be accessing enough resources in the evening to avoid torpor except in emergencies, while for smaller hummingbirds that cannot routinely store enough energy to avoid torpor at night, the routine hypothesis could perhaps better explain their torpor use patterns (though migrating birds offer exceptions to this trend; S. Hiebert, 1993). These results are consistent with previous findings that some combination of resource availability and energy storage affects torpor use (Powers et al., 2003), but also that temperature is not the main factor in determining whether an individual enters torpor or not—highlighting the importance of evaluating torpor under conditions of an organism’s natural environment.

Among the birds that used torpor, torpor bout duration was influenced more by night length than by chamber temperature, as predicted in the circadian rhythm hypothesis (Carpenter, 1974). Given that hummingbirds have limited energy stores, we expected colder nights to result in longer torpor bouts, but this was not reflected in torpor duration. Torpor bout duration was slightly shorter at the temperate sites than at the tropical sites, perhaps reflecting the shorter temperate nights. Although duration varied between sites, the proportion of the night spent in torpor was similar in temperate and tropical birds, indicating the importance of night length as a predictor of torpor duration. To confirm that torpor duration is affected by night length, additional torpor duration measurements are required for the same species either at different times of the year at temperate sites, or across latitudes with different day lengths at the same time of year.

Hourly energy savings were higher at colder, temperate sites than at warmer, tropical sites and were a function of how close T_a_ was to the bird’s minimum T_b_. There was no correlation between energy savings and torpor bout duration (Supplement S1: Fig. S9b). The tropical green-crowned brilliant saved slightly more energy per hour of torpor at the slightly colder Santa Lucia site, where it presumably approached minimum T_b_, than at the warmer Maquipucuna site. We did not measure minimum T_b_ for this species, but the T_a_ at these sites (average 25°C) was likely warmer than its minimum T_b_ because the highest reliable reported minimum T_b_ for hummingbirds is 18°C (Bech et al., 1997; Hainsworth et al., 1977), and our data show these birds’ torpid energy expenditures continued to decrease as nighttime temperatures got colder. The temperate broad-billed hummingbird saved more energy per hour of torpor at the warmer Sonoita site than at the colder Harshaw site (Fig. 3,5). Harshaw reached T_a_ and T_c_ below the broad-billed hummingbirds’ minimum T_b_, (∼15°C, Figs. 2, 3, 5), while Sonoita did not, allowing for maximal torpid energy savings at Sonoita and sub-maximal savings at Harshaw because the birds increased their metabolic rate to maintain their minimum T_b_.

**Figure 5:**
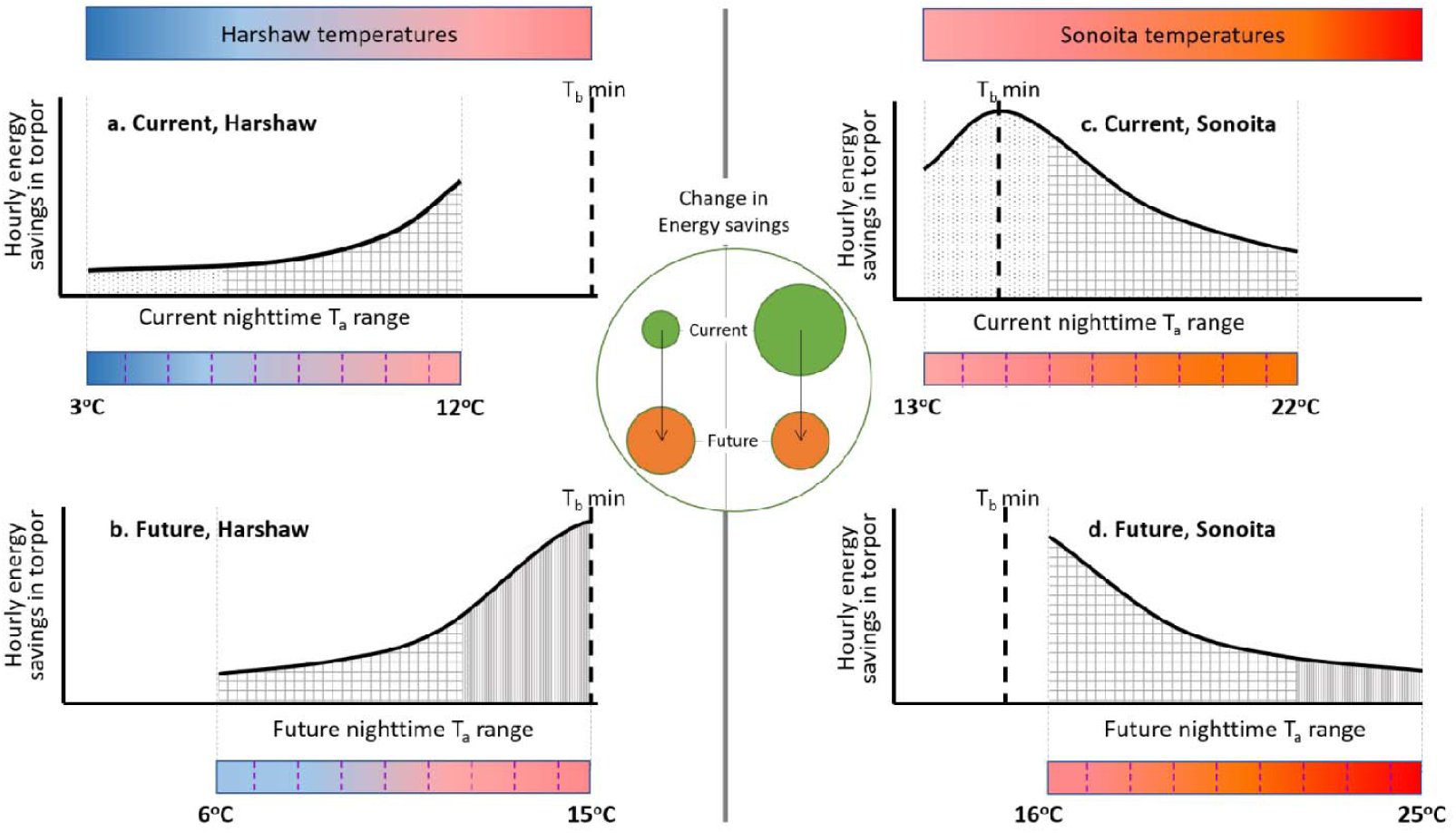
Schematic diagram depicting the relationship between hourly energy savings (calculated as % energy saved/hour of torpor relative to normothermy), minimum Tb, and Ta for the broad-billed hummingbird under current and future temperatures at two Arizona sites (Harshaw and Sonoita). Overall, energy savings depend on how close T_a_ is to minimum T_b_. Assuming a future increase of 3°C in nighttime temperatures, energy savings could decrease in Sonoita and increase in Harshaw under warming conditions. Color bars and temperature scales at the base of each plot represent temperature ranges at that time period and site. Minimum T_b_ for broad-billed hummingbirds (∼15°C) is depicted by the bold vertical dashed lines. Lighter vertical dashed lines represent the range of ambient temperatures for that time period. The ‘current’ plots have light dotted shading; future portions of the plots have dense vertical shading; portions of the plots that overlap have checkered shading. The circle in the middle represents overall nighttime energy savings under that scenario-green is current and orange is future.

In hummingbirds, a few studies have reported minimum T_b_ ranging from 7-18°C (Carpenter, 1974; Hainsworth & Wolf, 1970), while others mention that torpor savings are maximized at T_a_’s of 15-20°C (Kruger et al., 1982); however, the link between energy savings and the difference between T_a_ and T_b_ has not been explicitly articulated. Our results are consistent with some laboratory studies on mammals and birds (Hainsworth & Wolf, 1970; Jastroch et al., 2016), which have found that energy savings in torpor are maximized when T_a_ is approximately equal to minimum T_b_.

## CONCLUSION

Taking all its components into account, nighttime energy expenditure was affected most by torpor duration rather than by differences in hourly energy savings (Fig. 4). Hourly energy savings varied significantly between temperate and tropical sites, but the range of variation in energy savings did not have a significant effect on nighttime energy expenditure. Together, our findings and past literature (Powers et al., 2003) suggest that a hummingbird is likely to enter torpor on a given night if its stored energy drops below some threshold, and once it enters torpor, it maximizes torpor bout duration (Fig. 4). The energy a hummingbird saves per hour of torpor is negatively related to T_a_ – T_b_ min. On one hand, warming temperatures could increase energy savings from torpor for hummingbirds at very cold sites like Harshaw because T_a_-T_b_ would be reduced. On the other hand, warming temperatures could diminish energy savings from torpor at warmer tropical sites, like Maquipucuna and Santa Lucia or in moderate temperate environments like Sonoita, that currently have nighttime temperatures around minimum T_b_ (Fig. 5). However, warming temperatures at all our sites (where nights are colder than normothermic thermoneutral temperatures) would allow hummingbirds to lower energy expenditure during normothermy and cause an overall decrease in nighttime energy expenditure (Supplement S1: Table S4). As long as torpor can be used for long durations (contingent on night length), rising temperatures will likely not have a significant effect on total nighttime energy expenditure. Warming will likely have a larger effect on energy expenditure during nighttime hours spent in normothermy than during hours spent torpid (Supplement S1: Table S4).

## DATA ACCESSIBILITY

The datasets supporting this article have been uploaded as part of the supplementary material. Code used to run analyses (on R version 3.4.4) in this article are available at https://github.com/nushiamme/Torpor.

### ACKNOWLEDGEMENTS

We thank JL Andrew, NM Camacho, JR Canepa, KM Langland, SB Nutter and BG Weinstein for help with data collection. LM Dávalos, JS Levinton, HJ Lynch, P Copenhaver-Parry, and LR Yohe for helpful comments on the manuscript or statistical design. Maquipucuna and Santa Lucia Ecolodges, Arizona State Parks, the Southwestern Research Station, and the Hummingbird Monitoring Network provided logistical support. NASA (grant NNX11AO28G to CHG, Scott J. Goetz, SMW, and DRP), the Tinker Foundation, National Geographic Society (grant 9506-14 to AS), Stony Brook Department of Ecology and Evolution (to AS), a George Fox University Faculty Development Grant (GFU2014G02 to DRP), and the George Fox University Richter Science Scholar Grant (to RJS) funded the project. The project was approved by IACUC of Stony Brook University (IRBNet number: 282617-6), US Fish and Wildlife in Arizona (USFWS MB75714A-0), and the Ministry of Environment in Ecuador (Scientific investigation number 17-2014-IC-FAU-DPAP-MA).

## AUTHOR CONTRIBUTIONS

All authors were involved with study conception and design. AS, RS and DRP collected the data; AS and RS processed and analysed data and drafted the manuscript. CHG and DRP provided major comments and revisions. SMW provided comments on the manuscript. All authors gave final approval for publication.

